# IL4Pred2: Prediction of Interleukin-4 Inducing Peptides in Human and Mouse

**DOI:** 10.1101/2025.04.23.650150

**Authors:** Ritu Tomer, Naman Kumar Mehta, Shivani Malik, Shipra Jain, Gajendra P. S. Raghava

## Abstract

In 2013, our group developed IL4pred, a host-independent method for predicting interleukin-4 (IL-4) inducing peptides, which has been widely used by the scientific community. In this study, we present a second-generation method, IL4Pred2, which is a host-specific approach designed to predict IL-4 inducing peptides separately for human and mouse hosts. All models were trained, tested, and benchmarked on experimentally validated data obtained from the IEDB. We employed a wide range of state-of-the-art techniques for prediction, including similarity-based approaches, machine learning, deep learning methods, and large language models. Our best model achieved highest AUC 0.80 with MCC 0.45 for human and AUC 0.82 with MCC 0.50 for mouse on independent set of main datasets. All models were trained, test and optimized on training dataset. We validate our final model on an independent dataset which is not used in training or hyperparameter optimization of models. In this study, we developed models on three types of datasets called Main, Alternate1 and Alternate2 for predicting IL-4 inducing peptides. This abstract show performance of our models on Main dataset, performance of models on other datasets have been discussed in manuscript. One of the major objectives of this study is to facilitate research community in the area of immunotherapy and vaccine development. Thus, we developed, a web server and standalone software IL4pred2 for predicting, designing and scanning IL-4 inducing peptides in proteins (https://webs.iiitd.edu.in/raghava/il4pred2/ and https://github.com/raghavagps/il4pred2).

## Introduction

Interleukin-4 (IL-4) plays a critical role in various biological and immunoregulatory functions. It regulates a wide range of cells that include B-/T- lymphocytes, NK cells, monocytes, dendritic cells, basophils, mast cells, and fibroblasts [1–5]. It drives the differentiation of T-helper cells and promotes the growth of CD8+ T cells [6]. Its receptors appear on many cell types, primarily of hematopoietic origin [7, 8]. Researchers also refer to IL-4 as B cell growth factor-1 (BCGF-1) or B cell stimulatory factor-1 (BSF-1) [5, 9]. It also regulates the growth and differentiation of hematopoietic stem cells [1]. IL-4 activates the JAK/STAT signaling cascade, which leads to inflammation and hypersensitivity [10]. As illustrated in Figure 1, IL-4 is a key mediator of type 2 immunity. Recent studies have identified its therapeutic role in multiple myeloma, cancer, psoriasis, mucormycosis, and arthritis [11–14]. Current therapies for asthma and atopic dermatitis work by blocking IL-4 activity. In contrast, IL-4 has shown therapeutic benefits in animal models of spinal cord injury, stroke, and multiple sclerosis [15].

**Fig 1:**
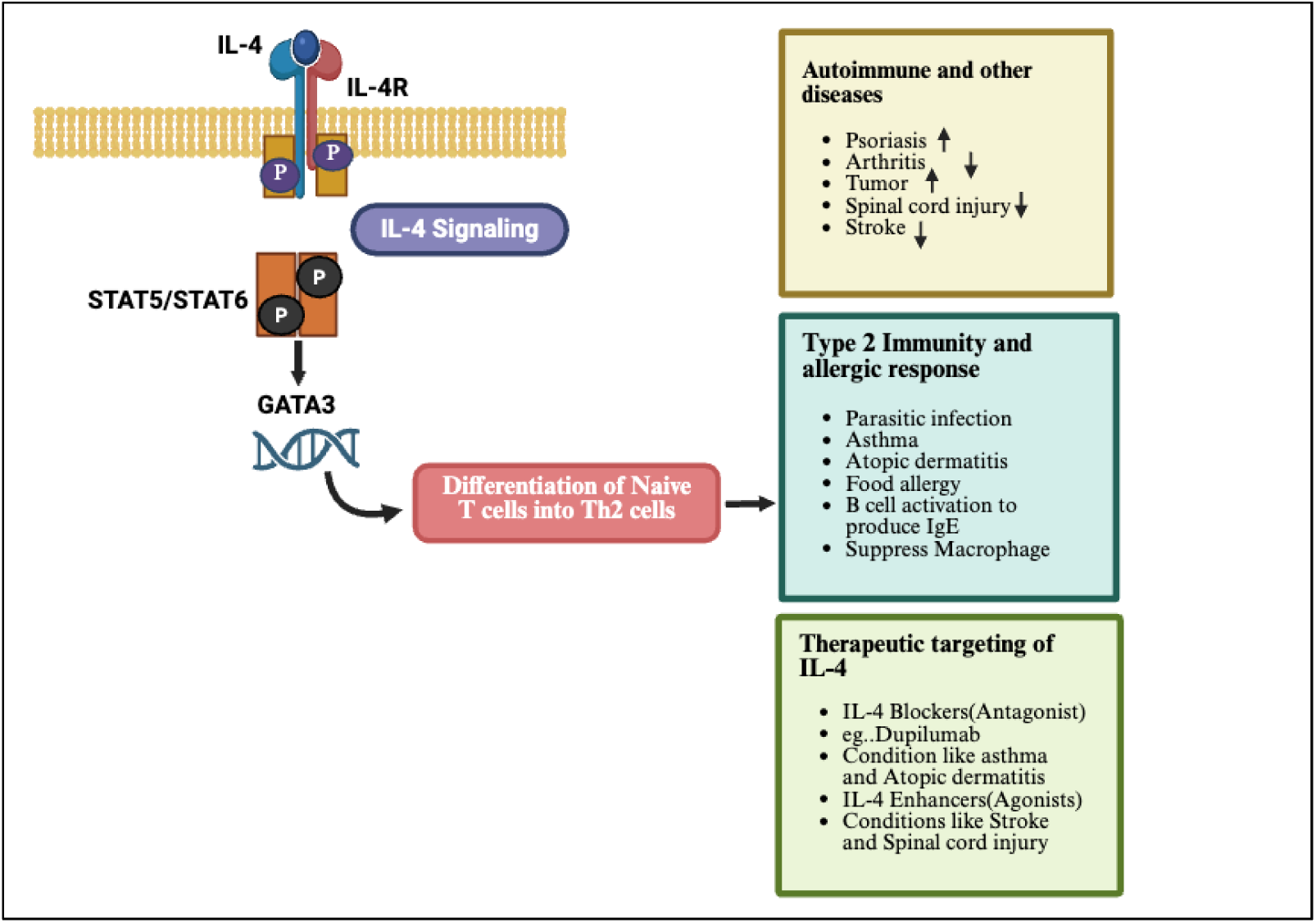
Schematic representation of role of IL4 cytokine in biological systems.

**Fig 2:**
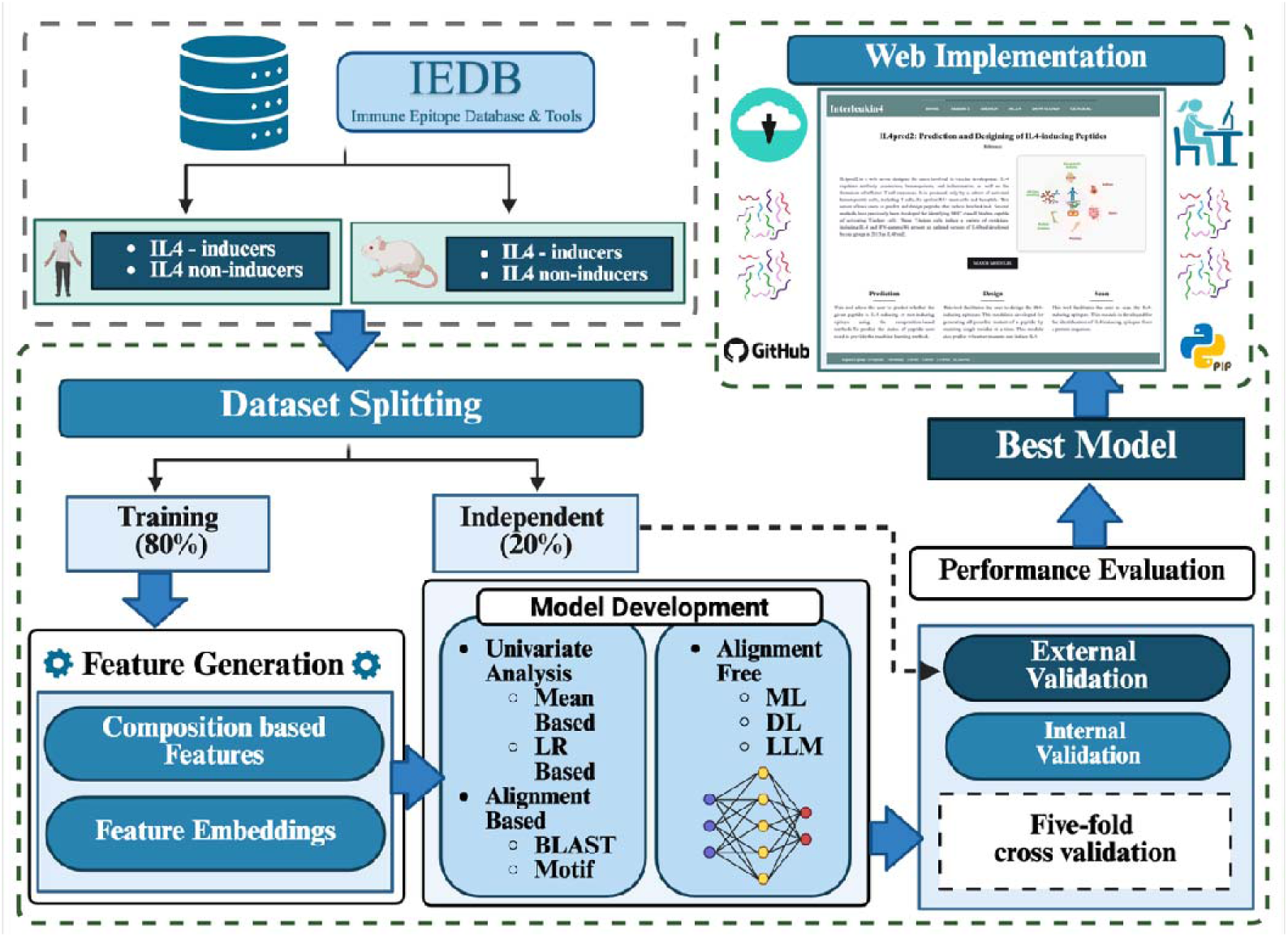
The figure depicts the overall architecture of the study.

Identifying peptides or antigenic regions capable of inducing IL-4 is, therefore, crucial for designing IL-4-based immunotherapies. Although experimental methods can detect IL-4 inducing peptides with high precision, they remain expensive, labor-intensive, and time-consuming. This highlights the need to develop efficient computational approaches for predicting and designing IL-4 inducing peptides. Over the years, researchers have developed several computational methods to predict cytokine-inducing peptides for a wide range of cytokines, which include IL-6, IL-7, IL-10, IL-13, interferon-gamma, and TNF-alpha [16–21]. In 2013, our group was the first to introduce IL4Pred, a method for predicting IL-4 inducing peptides [22]. IL4Pred has been widely used and cited by researchers to identify and experimentally validate IL-4 inducing peptides. In the past decade, several new methods have emerged for IL-4 prediction, including Meta-IL4 and PLM-IL4 [6, 23].

One of the major limitations of existing methods is their reliance on limited data available at the time of development. Another key drawback is their host-independent nature; these methods use a single model for all hosts. These method assuming all methods have same immune response for an antigen, which is not universally true. To address these limitations, we developed a second-generation method, IL4Pred2, which builds separate models for human and mouse to predict IL-4 inducing peptides. In addition, datasets used in this study have maximum and latest information from available sources. In this study, we retrieved experimentally validated peptides from the IEDB database to construct our datasets. Generating a positive set of IL-4 inducing peptides was straightforward, as such data are readily available in IEDB. However, constructing an appropriate negative dataset that is acceptable to the scientific community remains a significant challenge. To overcome this, we created three distinct datasets for each host, termed *Main*, *Alternate1*, and *Alternate2*. Each dataset includes a positive set of experimentally validated IL-4 inducing MHC class II binders; 845 for human and 560 for mouse. The negative set in each case contains an equal number of peptides that are not known to induce IL-4. Specifically, the Main dataset includes a mixture of MHC class II binders and non-binders as negatives; Alternate1 uses only MHC class II binders; and Alternate2 uses only MHC class II non-binders. Further details are provided in the Materials and Methods section, while the rationale for using multiple datasets is discussed in the Discussion section. We developed a wide range of models for each dataset and for both human and mouse hosts. These models are intended to aid in identifying potential antigens for immunotherapy and vaccine design [24].

## Materials & Methods

In this study, we made an effort to classify IL4-inducing peptides based on different hosts for which we have followed the below given methodology.

### 1.1. Dataset and Preprocessing

In this study, we developed models for human and mouse host, thus we create separate datasets for each host. We extract data from IEDB for creating datasets, IEDB contain experimentally validated peptides. For each host, we created three datasets termed as Main, Alternate1 and Alternate2 (**Please refer Supplementary Figure S1**). In all three datasets, positive dataset is same that is IL-4 inducing MHC II binding peptides (**Table 1**). We have taken different type of data to create negative dataset in three datasets. The negative dataset of Main dataset contains MHC Class II binders and MHC non-binders. We also make sure that none of the peptide in negative dataset have ability to induce IL-4. In case of Alternate1 dataset, our negative contains MHC Class II binders which do not induce IL-4. In case of Alternate2 dataset, our dataset contains MHC Class II non-binders which does not induce IL-4.

**Table 1:** The table shows the number of peptides in different datasets for human and mouse.

| Human |  |  |  |  |  |  |
| --- | --- | --- | --- | --- | --- | --- |
| Data Type | Main Dataset |  | Alternate1 Dataset |  | Alternate2 Dataset |  |
|  | Positive | Negative | Positive | Negative | Positive | Negative |
| Unique peptide | 900 | 1389 | 900 | 534 | 900 | 1855 |
| Length (9-22) | 845 | 1333 | 845 | 516 | 845 | 1817 |
| Final Data | 845 | 845 | 845 | 516 | 845 | 845 |
| Mouse |  |  |  |  |  |  |
| Data Type | Main Dataset |  | Alternate1 Dataset |  | Alternate2 Dataset |  |
|  | Positive | Negative | Positive | Negative | Positive | Negative |
| Unique peptide | 660 | 1787 | 660 | 905 | 660 | 882 |
| Length (9-22) | 560 | 1547 | 560 | 698 | 560 | 849 |
| Final Data | 560 | 560 | 560 | 560 | 560 | 560 |

We have selected MHC-class II non-binders by considering there IC_50_ values. As given in previous studies, MHC class II binding affinities were categorized based on their IC_50_ values, as good binders (IC_50_ ≤ 1000 nM), weak binders (IC_50_ 1,000– 10,000 nM), and non-binders (IC_50_ > 10000 nM) [25, 26]. In present study, we have flagged the peptides with IC_50_ value more than 1000 nM as MHC class II non-binders. Further, we preprocessed extracted data by removing redundant peptide sequences and applying length filter. We choose the length range based on maximum sequence coverage. The detailed data statistics of all three models for both hosts can be seen in **Table 1**.

### 1.2. Feature Extraction

We have deployed Pfeature tool to extract the peptide-based features from the dataset sequences [27]. Using this method, we have generated a total of 9189 feature vectors for every given sequence of our data. In addition to this, we have also computed composition-based features separately such as AAC, DPC and TPC to develop machine learning based models over single composition-based features.

### 1.3. Alignment based prediction methods

#### a) BLAST

BLAST tool is widely used for annotating and aligning peptide/ protein sequences [28–30]. In this study, we have implemented this tool to identify similarity of a query sequence with IL4 inducing peptides by aligning it to a manually curated database of IL4 inducing and non-inducing interleukins sequences. We deployed the BLASTshortP (version 2.9.0), a method designed for annotating short peptides, based on the resemblance among the sequences. In order to run this tool, we first created a repository of IL4 inducing and non-inducing peptides of training dataset. Then, each sequence in the validation dataset was searched against it. In this approach, we employed the top BLAST hit approach. In this method, a peptide sequence was classified as an inducing or non-inducing peptide based on the top match peptide label, using various E-value cutoffs ranging from 10^−1^ to 10^−6^. For example, if the query sequence hits a IL4 inducing peptide, then the sequence was categorized as IL4 inducing, otherwise, vice versa. The performance was evaluated using different E-value cutoffs.

#### b) Identification of Motifs

In the present study, we scanned exclusive conserved patterns in positive and negative peptide sequences using MERCI – Motif EmeRging and with Classes Identification and MEME (Multiple Em for Motif Elicitation) suits tools such as MEME-MAST (Motif Alignment and Search Tool) [31–33]. These two are widely used tools for motif discovery in each set of sequences. These tools aid in discovering exclusive patterns/ motifs and their occurrences that can distinguish between IL4 inducing and non-inducing peptides.

### 1.4. Alignment-free prediction methods

#### a) Machine learning models

With the aim to screening the IL4 inducing and non-inducing peptides, we deployed various widely used machine learning models such as Decision Tree, Random Forest, Logistic Regression, XGBoost (Extreme Gradient Boosting), K-Nearest Neighbors, Gaussian Naive Bayes, Extra Tree and Support Vector Classifier [21, 34]. To implement these machine learning algorithms, we deployed sklearn, a python-based machine learning library.

#### b) Feature Selection

In this study, we applied several widely known feature selection techniques such as Support Vector Classifier with L1 regularization (SVC-L1), Minimum Redundancy Maximum Relevance (mRMR), and Sequential Feature Selection (SFS). SVC-L1 is a type of Support Vector Machine with linear kernel, penalized with L1 regularization (also known as Lasso regularization) [35]. MRMR aims to extract features that have high relevance to the target variable with minimum redundancy with each other [36]. SFS is based on a greedy search algorithm that selects features sequentially [37].

#### c) Statistical method for feature reduction

In addition to the popular feature selection techniques, we have also applied mean based univariate analysis to select relevant features. In this step we have implemented the mean based univariate approach, by applying a student T-test. It enables us to compare the mean difference of the feature set between the inducing and non-inducing peptides. We ranked the features with significant p-values (p<0.05) and higher mean difference among both classes. Using these ranked features, we developed ML models using top 100, 150, 200, 250 feature sets.

#### d) Deep learning models

In present study, we have implemented 1D Convolutional Neural Networks (1D CNN) and TabNet for screening of IL4 inducing peptides. The 1D CNN extracts spatial patterns from peptide sequences encoded using one-hot representation [38, 39]. TabNet, designed for tabular data, leverages feature transformer blocks to model complex relationships and attentive transformers for feature selection, ensuring interpretability through sparse regularization [40]. While 1D CNN processes sequence-based information, TabNet focuses on tabular features like physicochemical properties.

#### e) Reinforcement learning

In this study, we have also implemented the LLM (large language model) for the classification of IL4 inducing and non-inducing peptides. We utilized ProtBERT which is a transformed based Bert model and has been pretrained on a large pool of protein sequences in a self-supervised fashion [41]. We extracted embeddings from the pretrained model, and which represented the amino acid sequences. These embeddings encode biochemical and structural features of peptides, such as similarities, motifs, and functional relationships, learned during pretraining on large protein sequence datasets. Post that we fine-tuned the ProtBERT model on a labelled dataset and varied epoch values (3 to 7) to record the performance parameters.

### 1.5. Cross validation

In order to maintain the machine learning standards, we randomly split all datasets created as 80% training data and 20% independent validation dataset. In addition to this, we have implemented a 5-fold cross validation technique over our training dataset, to ensure robustness and to prevent overfitting of our model. Using this technique, we have divided our training data (i.e. 80% of dataset) into five subsets, in which four are used as training and one as testing dataset [18, 21, 42]. This process is iterated five times so that each subset is used as a testing subset. Post that the best model is implemented on an independent validation set (i.e. 20% of dataset) for evaluating the machine learning based classification model performance.

### 1.6. Performance evaluation

The efficacy of our machine learning based prediction models developed for the prediction of IL4 inducing and non-inducing peptides, we calculated widely used evaluation parameters in literature [22, 43]. In the present study, we implemented both threshold-dependent parameters such as sensitivity (Sens), specificity (Spec) and accuracy (Acc) and independent parameters such as Area Under the Receiver Operating Characteristic (AUROC) curve to measure the performance of the models. These parameters were calculated using the following equations:

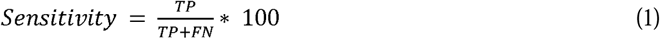

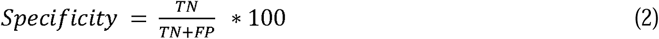

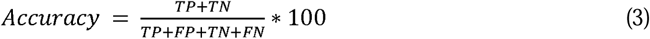

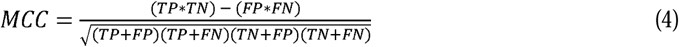

Where, FP is false positive, FN is false negative, TP is true positive, and TN is true negative.

### 1.7. Ensemble approach

In present study, we have developed an ensemble method that integrates machine learning models with traditional sequence-based approaches to enhance predictive performance. Specifically, we combined the top-performing machine learning models with alignment-based techniques, such as BLAST, as well as motif-based methods, including MEME-MAST and MERCI. By leveraging the complementary nature of these approaches, we intend to capture evolutionary relationships through sequence alignment and functional motifs through pattern recognition, ultimately leading to combined prediction model. Using this we developed ensemble approach using 3 methods: 1) BLAST mapped IL4 inducing peptides based on similarity hits then ML model was used to predict the non-mapped peptides. 2) MERCI based motif sequence patterns were identified in the validation dataset and then, ML model predicted the non-identified peptides. 3) Similarly, combined MEME MAST motifs with ML model prediction results.

## Results

### 1. Preliminary analysis

#### 1.1. Positional Analysis

In this study, we did positional analysis to understand the amino acids preferences at specific positions in our main dataset between IL4-inducing and IL4 non-inducing peptides for human and mouse dataset. Two sample logo positional analysis depicts the dominance and relative occurrence of a particular amino acid in the sequence [44]. It is vital to note that the first eight positions in the logo depicts the eight N-terminal residues, and the last eight depicts the eight C-terminal residues. It is evident from **Figure 3.1**, that in human host lysine (K), isoleucine (I) and tyrosine (Y) are found to be predominant at multiple positions in IL4 inducing peptides. Lysine is often present at 1^st^, 4^th^, 5^th^, 8^th^, 9^th^, 10^th^,13^th^ and 18^th^ position. Whereas proline (P), glycine (G), leucine (L) and aspartic acid (D) amino acids are in abundance in the non-inducing peptides. Proline is often present at 1^st^, 5^th^, 7^th^, 8^th^, 9^th^, 13^th^, 15^th^ and 18^th^ position.

**Fig 3.1:**
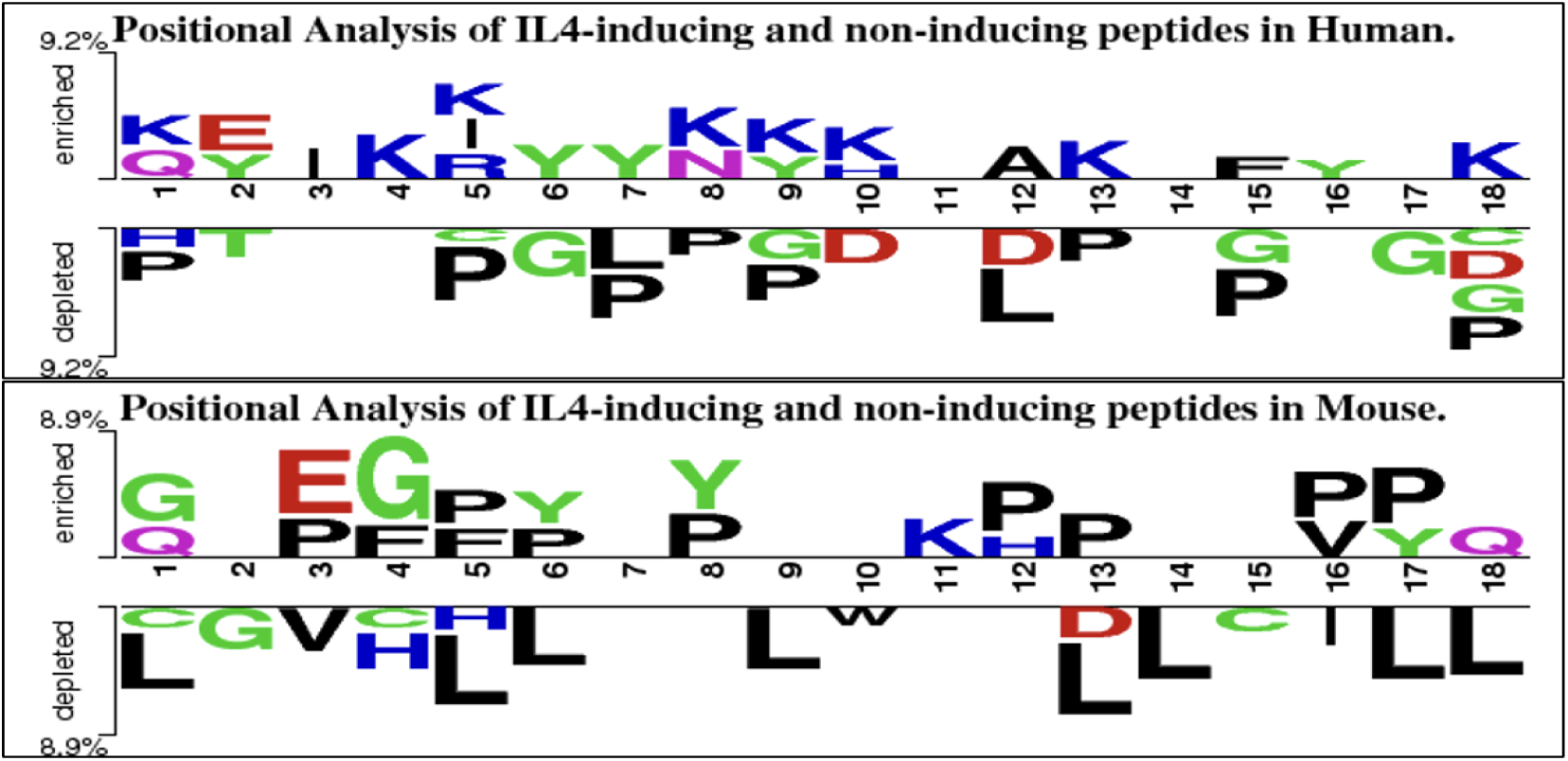
This figure represents two-sample logo positional analysis for IL4 Inducing and Non-inducing peptides in human and mouse host.

**Fig 3.2:**
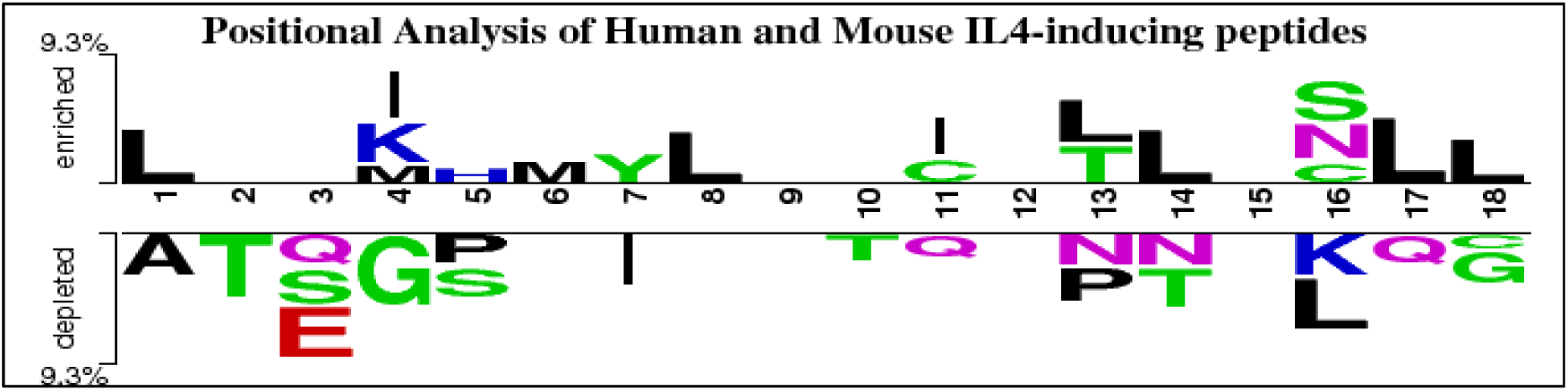
This figure represents two-sample logo positional analysis between IL4 Inducing peptides of human and mouse host.

In contrast, in mouse host certain amino acids are found to be confined at a particular position. In this data, glycine (G) at 1^st^ and 4^th^, proline (P) at 3^rd^, 5^th^, 6^th^, 8^th^, 12^th^, 13^th^, 16^th^ and 17^th^, phenylalanine (F) at 4^th^ and 5^th^ and tyrosine (Y) are found to be present at 6^th^, 8^th^ and 17^th^ position in IL4 inducing peptides, and leucine (L) is found to be prominent at 1^st^, 5^th^, 6^th^, 9^th^, 13^th^, 14^th^, 17^th^ and 18^th^ position in IL4 non-inducing peptides.

We have also performed positional analysis to understand the amino acids preferences at specific positions in IL4-inducing peptides of human and mouse. Here, we observed that amino acids leucine (L) is abundant at 1^st^, 8^th^, 13^th^, 14^th^, 17^th^ and 18^th^ position, methionine (M) is found at 4^th^ AND 6^th^ position and cysteine is present at 11^th^ and 16^th^ position in human IL4-inducers while serine (S), glutamine (Q), proline (P), asparagine (N) and glycine (G) residues are abundant at specific positions in mouse IL4-inducing peptides.

#### 1.2. Compositional Analysis

In present study, we performed a compositional analysis of peptide dataset to capture the amino acid composition molecular insights as IL4 inducing peptides. We observed that in humans’ Glutamic acid, Isoleucine, Lysine and Tyrosine amino acid residues were found in abundance in IL4 inducing peptides, whereas Aspartic acid, phenylalanine, histidine, Leucine, Proline, Asparagine and Threonine were reported in higher frequency in non-inducing peptides. As depicted in **Figure 4.1**, amino acid composition trends in mouse host were not like humans. In mouse, Proline, Serine and Tyrosine average amino acid composition were higher in IL4 inducing peptides, whereas Cysteine, Histidine, Isoleucine, Leucine, Methionine and Arginine are on abundant side in non-inducing peptides.

**Figure 4.1:**
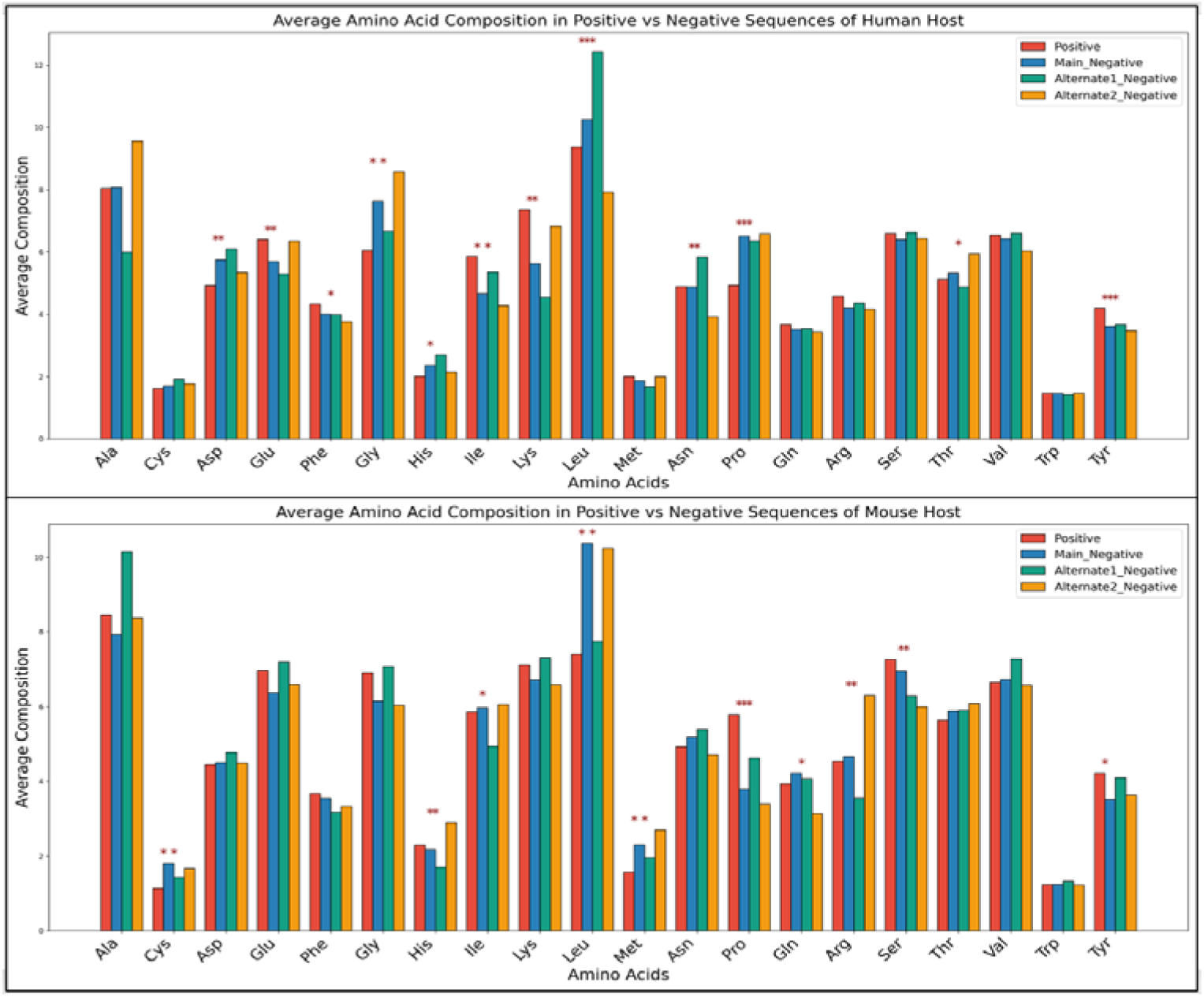
This figure depicts IL4 inducing and non-inducing peptides amino acid composition in human and mouse hosts.

**Figure 4.2:**
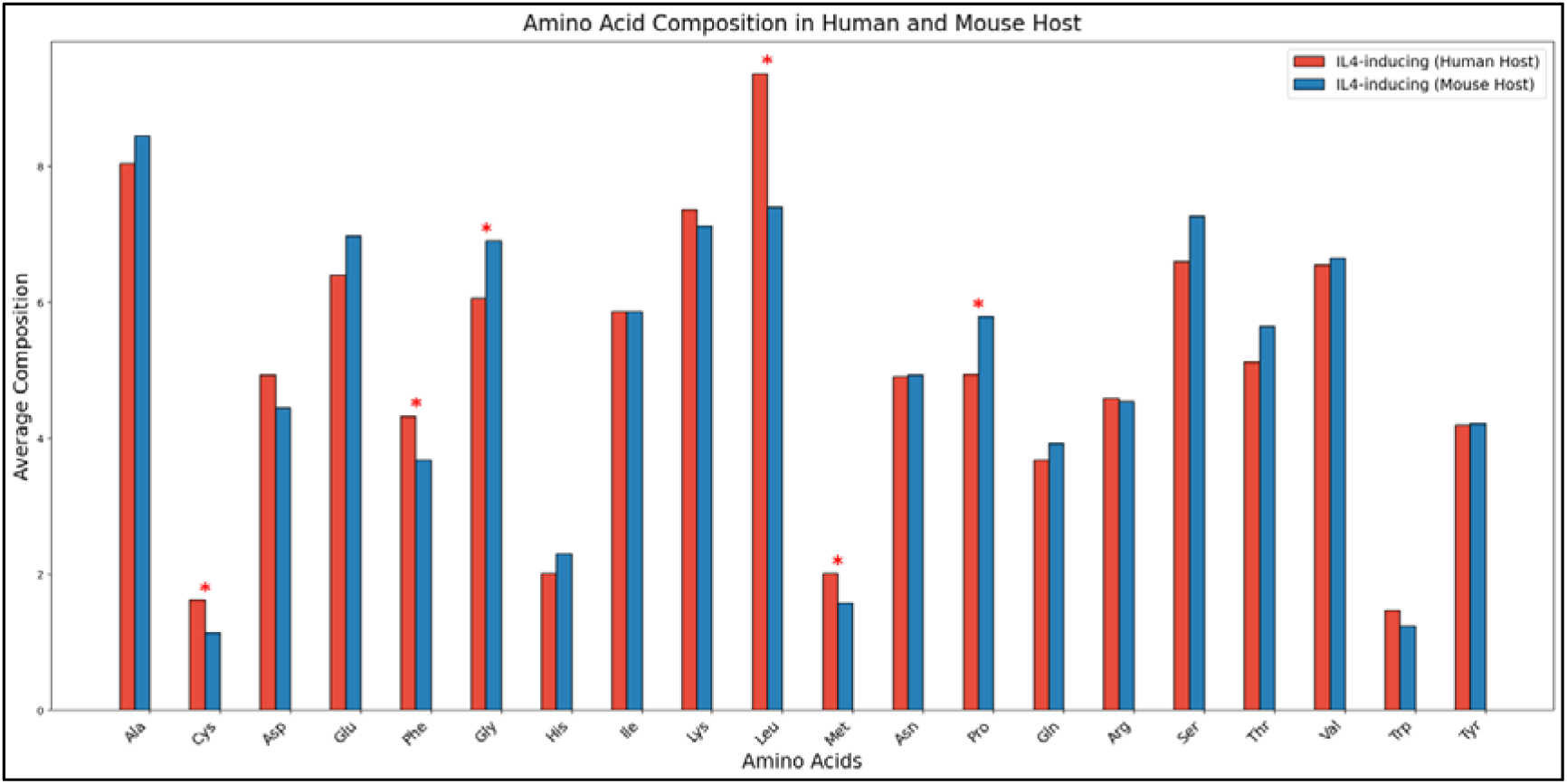
This figure shows amino acid composition of IL-4 inducing peptides in human and mouse.

To understand the difference between human and mouse specific IL4 inducing peptides, we performed average compositional analysis between IL4 inducing peptides of human and mouse hosts. As we hypothesized that there is difference between composition of human and mouse data. Here, we can clearly see that the average amino acid composition of cysteine, phenylalanine, leucine and methionine residues are significantly higher in human IL4-inducing peptides while average amino acid composition of glycine and proline residues are abundant in mouse data. The same trend can also be seen in amino acid positional analysis between human and mouse IL4-inducing peptides.

#### 1.3. Mean based analysis

In order to capture the mean-based differentiation among feature pool, we ranked them based on their higher mean difference between IL4 inducing and non-inducing peptides with significant p-value. As observed from **Table 2** that in human host, Pseudo Amino Acid Composition (PAAC) feature emerged on top with a difference of 0.34, whereas composition enhanced transition and distribution (CeTD) based features appeared on top with difference of 0.36 in mouse dataset. The complete list of features with significant p-value and their ranks can be referred in **Supplementary Table S1**.

**Table 2:** The table depicts top 5 features which have maximum difference in positive and negative peptides.

| Main Dataset of Human |  |  |  | Main Dataset of Mouse |  |  |  |
| --- | --- | --- | --- | --- | --- | --- | --- |
| Feature | Pos_avg | Neg_avg | Diff | Feature | Pos_avg | Neg_avg | Diff |
| PAAC1_lam1 | 0.17 | -0.17 | 0.34 | CeTD_100_p_VW1 | 0.18 | -0.18 | 0.36 |
| SOC1_SC1 | 0.16 | -0.16 | 0.32 | CeTD_50_p_VW1 | 0.18 | -0.18 | 0.36 |
| CeTD_12_PO | -0.15 | 0.15 | -0.30 | SER_P | -0.17 | 0.17 | -0.34 |
| SER_K | -0.14 | 0.14 | -0.28 | PAAC1_L | -0.17 | 0.17 | -0.34 |
| QSO1_G_P | -0.14 | 0.14 | -0.28 | APAAC1_L | -0.17 | 0.17 | -0.34 |

**Note:** Diff: Difference between Positive mean and Negative Mean;

#### 1.4. Single based feature performance

In present study, we performed Logistic regression-based analysis to understand the efficacy of single feature to discriminate IL4 inducing and non-inducing peptides. As evident from **Table 3**, we observed that PAAC1_lam1 feature reported the maximum AUC as 0.59 for human host. Few of these features were also observed in the top 5 list of highest significant mean difference list. Whereas composition enhanced transition and distribution-based features (CeTD_100_p_VW1, CeTD_50_p_VW1) reported the maximum AUC as 0.61 and 0.60 respectively for the mouse host which are also observed in the list of top 5 highly significant mean difference list. Please refer **Supplementary Table S2** for the detailed list.

**Table 3:**
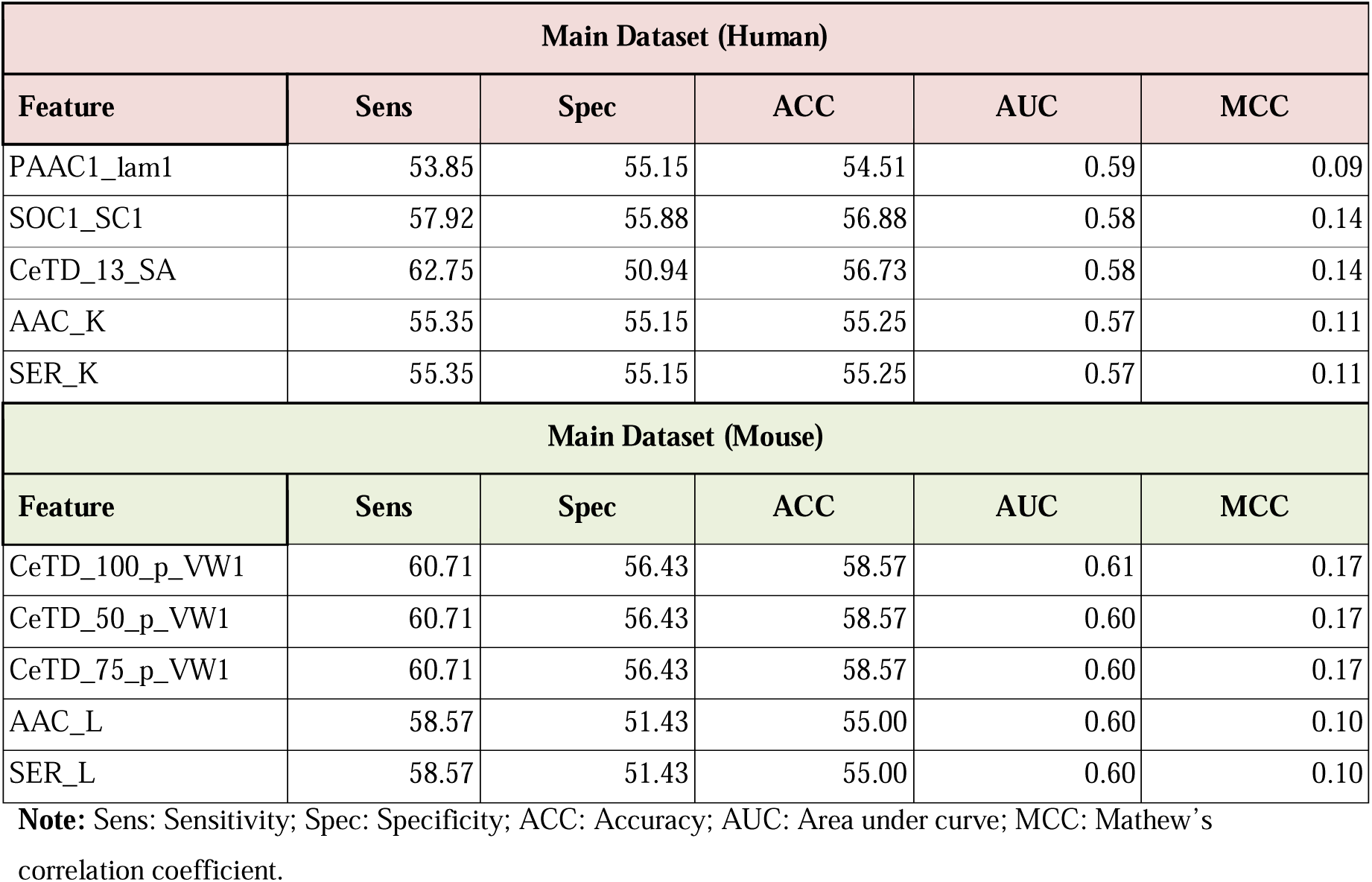
Single feature-based performance of top 5 features where models were developed using logistic regression.

| Main Dataset (Human) |  |  |  |  |  |
| --- | --- | --- | --- | --- | --- |
| Feature | Sens | Spec | ACC | AUC | MCC |
| PAAC1_lam1 | 53.85 | 55.15 | 54.51 | 0.59 | 0.09 |
| SOC1_SC1 | 57.92 | 55.88 | 56.88 | 0.58 | 0.14 |
| CeTD_13_SA | 62.75 | 50.94 | 56.73 | 0.58 | 0.14 |
| AAC_K | 55.35 | 55.15 | 55.25 | 0.57 | 0.11 |
| SER_K | 55.35 | 55.15 | 55.25 | 0.57 | 0.11 |
| Main Dataset (Mouse) |  |  |  |  |  |
| Feature | Sens | Spec | ACC | AUC | MCC |
| CeTD_100_p_VW1 | 60.71 | 56.43 | 58.57 | 0.61 | 0.17 |
| CeTD_50_p_VW1 | 60.71 | 56.43 | 58.57 | 0.60 | 0.17 |
| CeTD_75_p_VW1 | 60.71 | 56.43 | 58.57 | 0.60 | 0.17 |
| AAC_L | 58.57 | 51.43 | 55.00 | 0.60 | 0.10 |
| SER_L | 58.57 | 51.43 | 55.00 | 0.60 | 0.10 |

**Note:** Sens: Sensitivity; Spec: Specificity; ACC: Accuracy; AUC: Area under curve; MCC: Mathew's correlation coefficient.

### 2. Alignment based prediction methods

Alignment-based methods rely on the similar patterns or presence of specific motifs in sequences. These types of methods have higher specificity but low sensitivity. In this study, we used following alignment or similarity-based methods.

#### 2.1. BLAST

In this approach, we preform BLAST search of peptides in independent dataset against peptides in training dataset. We mapped hits at several e-values ranging from 1e^−1^ to 1e^−6^. The query sequence was classified as IL4 inducer if its top hit was positive and as IL4 non-inducer if its top hit was from negative data. The detailed list of hits achieved by BLAST on human and mouse main validation data is shown below in **Table 4**.

**Table 4:** The performance of BLAST at different cut-off value.

| Human Host (Main Dataset) |  |  |  | Mouse Host (Main Dataset) |  |  |  |
| --- | --- | --- | --- | --- | --- | --- | --- |
| E-value | T_hits | CP_hits | CN_hits | E-value | T_hits | CP_hits | CN_hits |
| **1.00E-01** | 557 | 89 | 59 | **1.00E-01** | 197 | 30 | 29 |
| **1.00E-02** | 458 | 75 | 41 | **1.00E-02** | 164 | 28 | 27 |
| **1.00E-03** | 331 | 60 | 31 | **1.00E-03** | 126 | 22 | 20 |
| **1.00E-04** | 207 | 48 | 21 | **1.00E-04** | 80 | 14 | 12 |
| **1.00E-05** | 126 | 36 | 12 | **1.00E-05** | 39 | 9 | 6 |
| **1.00E-06** | 66 | 25 | 3 | **1.00E-06** | 18 | 7 | 1 |

Note: T\_hits: Total hits found; CP\_hits: correct positive hits; CN\_hits: correct negative hits.

#### 2.2. Motif analysis

We performed comprehensive motif analysis using two established tools—MERCI and MEME-MAST— to identify signature sequence patterns associated with IL-4 inducing and non-inducing peptides. Motifs were first extracted from the training dataset and then scanned across independent validation datasets from both humans and mouse to evaluate their presence and relevance. The top 10 motif hits identified by each tool, along with corresponding correct and incorrect classification outcomes, are summarized in **Table 5**. Detailed motif coverage and occurrence for the MERCI tool, along with ensemble method results, are provided in **Supplementary Table S3.1**. Similarly, comprehensive motif coverage and ensemble results for MEME-MAST analysis can be found in **Supplementary Table S3.2**.

**Table 5:** The list of Motif hits obtained using MERCI & MEME MAST on Human and Mouse validation datasets.

| Host | Classification Method | Number of K motifs | IL-4 inducing |  |  | IL-4 Non-inducing |  |  |  |
| --- | --- | --- | --- | --- | --- | --- | --- | --- | --- |
|  |  |  | motif | Correct | Incorrect | motif | Correct | Incorrect |  |
| MERCI |  |  |  |  |  |  |  |  |  |
| Human | None | 10 | 5 | 7 | 1 | 0 | 0 | 0 |  |
| Mouse | None | 10 | 1 | 1 | 0 | 3 | 6 | 3 |  |
| MEME-MAST |  |  |  |  |  |  |  |  |  |
| Host | E-value | Mini-site | Number of K motifs | IL-4 inducing |  |  | IL-4 Non-inducing |  |  |
|  |  |  |  | motif | Correct | Incorrect | motif | Correct | Incorrect |
| Human | 0.05 | 10 | 50 | 14 | 30 | 8 | 10 | 15 | 12 |
| Mouse | 0.05 | 10 | 10 | 4 | 6 | 3 | 7 | 5 | 2 |

### 3. Alignment free approach

#### 3.1. Machine learning based classifiers

In this study, we have applied several machine learning-based algorithms on composition-based features including amino acid composition (AAC), di-peptide composition (DPC) and tri-peptide composition (TPC). As evident in **Table 6**, for human host Extra tree classifier developed over AAC feature outperformed all and achieved 0.76 as AUC score. In addition to this, Random Forest classifier developed using DPC feature reported highest AUC as 0.70 with balanced sensitivity and specificity for mouse host. Detailed results on training and validation dataset can be referred in **Supplementary Table S4.**

**Table 6:** The performance of top machine learning based models developed using different type of compositional features on independent dataset.

| Human Dataset (Main data) |  |  |  |  |  |  |
| --- | --- | --- | --- | --- | --- | --- |
| Model | Feature | Sens | Spec | ACC | AUC | MCC |
| ET | AAC | 71.43 | 62.18 | 67.16 | 0.76 | 0.34 |
| ET | DPC | 70.33 | 66.67 | 68.64 | 0.76 | 0.37 |
| SVC | TPC | 65.39 | 67.31 | 66.27 | 0.74 | 0.33 |
| Mouse Dataset (Main data) |  |  |  |  |  |  |
| Model | Feature | Sens | Spec | ACC | AUC | MCC |
| RF | AAC | 64.57 | 61.86 | 63.39 | 0.68 | 0.26 |
| RF | DPC | 65.35 | 60.83 | 63.39 | 0.70 | 0.26 |
| SVC | TPC | 64.57 | 70.10 | 66.96 | 0.69 | 0.34 |

**Note:** Sens: Sensitivity; Spec: Specificity; ACC: Accuracy; AUC: Area under curve; MCC: Mathew's correlation coefficient.

#### 3.2. Feature Selection

In order to develop a robust model, we applied popular feature selection techniques like SVC-L1, mRMR and SFS techniques to extract important features from the 9189-feature pool generated using Pfeature. We selected 149 features from SVC-L1, 50 from mRMR and 17 from SFS techniques for human host. Similarly, we extracted 97 features from SVC-L1, 50 from mRMR and 17 from SFS techniques for mouse host. We developed machine learning classifiers using these selected features. As depicted in **Table 7**, we observed highest AUC as 0.79 using Extra Tree classifier developed SVC-L1 feature selection technique for human host and 0.72 using Random Forest classifier developed SVC-L1 feature selection technique for mouse host. Detailed performance evaluation metrics for training data can be referred in **Supplementary Table S5.**

**Table 7:** The performance of top machine learning based models developed on selected features obtained from different techniques.

| Method | Model | No of selected features | Human host |  |  |  |  |
| --- | --- | --- | --- | --- | --- | --- | --- |
|  |  |  | Sens | Spec | ACC | AUC | MCC |
| SVC-L1 | ET | 149 | 73.08 | 69.23 | 71.3 | 0.79 | 0.42 |
| mRMR | XGB | 50 | 67.03 | 56.41 | 62.13 | 0.68 | 0.24 |
| SFS | LR | 17 | 61.54 | 58.33 | 0.60 | 0.63 | 0.20 |
| Method | Model | No of features | Mouse host |  |  |  |  |
|  |  |  | Sens | Spec | ACC | AUC | MCC |
| SVC-L1 | RF | 116 | 69.29 | 60.83 | 65.63 | 0.72 | 0.30 |
| mRMR | XGB | 50 | 65.35 | 58.76 | 62.5 | 0.67 | 0.24 |
| SFS | LR | 17 | 56.69 | 64.95 | 0.60 | 0.60 | 0.21 |

**Note:** Sens: Sensitivity; Spec: Specificity; ACC: Accuracy; AUC: Area under curve; MCC: Mathew's correlation coefficient.

In addition to these feature selection techniques, we also performed the mean based univariate analysis to rank significant features among IL4 inducing and non-inducing peptides. The top ranked 100, 150, 200 and 300 features with significant p-values were selected and used to implement machine learning based classifiers. As evident in **Table 8**, we achieved a maximum AUC as 0.80 with MCC of 0.45 over top 300 features for human host while over top 300 features for mouse host we achieved AUC of 0.82 with MCC of 0.50 using Multilayer Perceptron (MLP) based machine learning classifier. See **Supplementary Table S6** for detailed results over training and validation set.

**Table 8:** The performance of top machine learning based models developed using different number of features selected using univariate analysis.

| Human host |  |  |  |  |  |  |
| --- | --- | --- | --- | --- | --- | --- |
| Feature | Model | Sens | Spec | ACC | AUC | MCC |
| Top_100 | ET | 0.71 | 0.71 | 71 | 0.76 | 0.41 |
| Top_150 | RF | 0.70 | 0.69 | 70 | 0.76 | 0.39 |
| Top_200 | ET | 0.71 | 0.68 | 70 | 0.76 | 0.39 |
| Top_250 | MLP | 0.75 | 0.69 | 72 | 0.76 | 0.44 |
| Top_300 | MLP | 0.75 | 0.7 | 73 | 0.80 | 0.45 |
| Mouse host |  |  |  |  |  |  |
| Feature | Model | Sens | Spec | ACC | AUC | MCC |
| Top_100 | GNB | 0.65 | 0.69 | 67 | 0.71 | 0.34 |
| **Top_150** | **GNB** | 0.63 | 0.72 | 67 | 0.72 | 0.35 |
| **Top_200** | **ET** | 0.72 | 0.65 | 69 | 0.73 | 0.37 |
| **Top_250** | **MLP** | 0.69 | 0.7 | 69 | 0.76 | 0.38 |
| **Top_300** | **MLP** | 0.74 | 0.76 | 75 | 0.82 | 0.50 |

**Note:** Sens: Sensitivity; Spec: Specificity; ACC: Accuracy; AUC: Area under curve; MCC: Mathew's correlation coefficient.

#### 3.3. Deep learning-based classifiers

In present study, deep learning-based classifiers have been deployed for the screening of IL4 inducing and non-inducing peptides. We have implemented 1D CNN and TabNet for classifying among both groups. As shown in **Table 9**, RRI (Repetitive Residue Information) feature reported the maximum AUC score as 0.66 for human host, whereas TPC feature achieved higher AUC score as 0.64 for mouse host using CNN classifier. **Supplementary Table S7** can be referred for performance score for training and testing dataset for both deep learning-based classifier methods.

**Table 9:** The performance of deep learning-based models developed using different types of residue compositions and evaluated on independent dataset (main dataset).

| Human host |  |  |  |  |  |  |
| --- | --- | --- | --- | --- | --- | --- |
| Model | Feature | Sens | Spec | ACC | AUC | MCC |
| AAC | CNNClassifier | 0.61 | 0.55 | 0.58 | 0.61 | 0.16 |
| DPC | CNNClassifier | 0.65 | 0.53 | 0.59 | 0.62 | 0.18 |
| DPC_LEN | CNNClassifier | 0.57 | 0.67 | 0.62 | 0.65 | 0.24 |
| RRI | CNNClassifier | 0.65 | 0.61 | 0.63 | 0.66 | 0.26 |
| TPC | CNNClassifier | 0.66 | 0.56 | 0.62 | 0.65 | 0.22 |
| AAC | TabNetClassifier | 0.44 | 0.6 | 0.51 | 0.51 | 0.04 |
| DPC | TabNetClassifier | 0.84 | 0.18 | 0.54 | 0.45 | 0.03 |
| DPC_LEN | TabNetClassifier | 0.35 | 0.74 | 0.53 | 0.58 | 0.09 |
| RRI | TabNetClassifier | 0.65 | 0.39 | 0.53 | 0.53 | 0.05 |
| TPC | TabNetClassifier | 0.03 | 0.95 | 0.45 | 0.53 | 0.06 |
| Mouse host |  |  |  |  |  |  |
| Model | Feature | Sens | Spec | ACC | AUC | MCC |
| AAC | CNNClassifier | 0.55 | 0.58 | 0.56 | 0.54 | 0.13 |
| DPC | CNNClassifier | 0.51 | 0.57 | 0.54 | 0.59 | 0.08 |
| DPC_LEN | CNNClassifier | 0.54 | 0.68 | 0.60 | 0.63 | 0.22 |
| RRI | CNNClassifier | 0.57 | 0.45 | 0.52 | 0.52 | 0.02 |
| TPC | CNNClassifier | 0.55 | 0.62 | 0.58 | 0.64 | 0.17 |
| AAC | TabNetClassifier | 0.50 | 0.58 | 0.54 | 0.56 | 0.08 |
| DPC | TabNetClassifier | 0.35 | 0.68 | 0.49 | 0.49 | 0.03 |
| DPC_LEN | TabNetClassifier | 0.01 | 0.98 | 0.43 | 0.51 | 0.05 |
| **RRI** | TabNetClassifier | 0.54 | 0.52 | 0.53 | 0.53 | 0.05 |
| **TPC** | TabNetClassifier | 0.01 | 0.98 | 0.43 | 0.51 | 0.05 |

**Note:** Sens: Sensitivity; Spec: Specificity; ACC: Accuracy; AUC: Area under curve; MCC: Mathew's correlation coefficient.

#### 3.4. Large Language model

In this study, we have implemented pretrained protBERT large language model for classification of IL4 inducing and non-inducing peptide groups. As shown in **Table 10**, we observed the maximum AUC as 0.72 using fine-tuned model while AUC of 0.75 achieved feature embeddings extracted from finetuned protBERT model for human host at an epoch value of 5. For the mouse host, the highest AUC of 0.66 was achieved at epoch 7 for both the fine-tuned model and feature embeddings. Refer **Supplementary Table S8** for performance matrix for all models.

**Table 10:** The performance of protein language model protBERT and machine learning models.

| Epochs | Feature | Model | Human host |  |  |  |  |
| --- | --- | --- | --- | --- | --- | --- | --- |
|  |  |  | Sens | Spec | ACC | AUC | MCC |
| 3 | Sequence | Finetuned | 0.79 | 0.50 | 65.98 | 0.72 | 0.31 |
|  | Embedding | RF | 0.72 | 0.69 | 71.00 | 0.75 | 0.41 |
| 5 | Sequence | Finetuned | 0.56 | 0.73 | 64.20 | 0.69 | 0.30 |
|  | Embedding | ET | 0.70 | 0.67 | 69.00 | 0.72 | 0.37 |
| 7 | Sequence | Finetuned | 0.65 | 0.69 | 67.46 | 0.70 | 0.35 |
|  | Embedding | ET | 0.69 | 0.63 | 66.00 | 0.71 | 0.32 |
| Epochs | Feature | Model | Mouse host |  |  |  |  |
|  |  |  | Sens | Spec | ACC | AUC | MCC |
| 3 | Sequence | Finetuned | 0.57 | 0.63 | 60.27 | 0.60 | 0.21 |
|  | Embedding | ET | 0.60 | 0.59 | 59.00 | 0.62 | 0.18 |
| 5 | Sequence | Finetuned | 0.69 | 0.52 | 62.05 | 0.63 | 0.22 |
|  | Embedding | ET | 0.64 | 0.60 | 62.00 | 0.64 | 0.23 |
| 7 | Sequence | Finetuned | 0.55 | 0.67 | 60.71 | 0.66 | 0.22 |
|  | Embedding | ET | 0.61 | 0.62 | 61.00 | 0.66 | 0.22 |

**Note:** Sens: Sensitivity; Spec: Specificity; ACC: Accuracy; AUC: Area under curve; MCC: Mathew's correlation coefficient.

#### 3.5. Ensemble/Hybrid Approach

##### 3.5.1. BLAST

For hybrid method, we merged the similarity-based prediction classifiers and machine learning algorithms. The BLAST hit results were merged with the best performing machine learning based classifier labels. We observed the maximum AUC score as 0.76 with MCC of 0.38 at an e-value of 1e^−6^ for with maximum correct hits for human host, whereas the highest AUC score reported in mouse host is 0.71 with MCC of 0.29 at an e-value of 1e^−6^. Detailed performance matrix can be referred in **Supplementary Table S9.**

##### 3.5.2. Motif

In this approach, we extracted exclusive motifs/ recurring patterns found in IL4 inducing and non-inducing peptides training dataset using MERCI & MEME tools. Then we searched for these patterns in independent validation dataset peptides and reported their presence. If a positive motif is found in query peptide, then it is marked as IL4 inducing peptide and +0.5 value is assigned and 0 if no match found. Finally, we combine predicted labels of best machine learning model with motif score. For MERCI tool we reported the maximum AUC as 0.80 with MCC of 0.45 for human host and AUC as 0.80 with MCC of 0.50 for mouse host by specifying maximum frequency of 10 positive motifs in MERCI tool using default parameter. Detailed motif coverage and occurrence list with ensemble method results for MERCI tool can be referred in **Supplementary Table S3.1**. Similarly, for MEME-MAST tool ensemble method we have achieved maximum AUC as 0.70 with MCC of 0.31 for human host and AUC as 0.70 with MCC of 0.28 for mouse host with p-value 0.05 respectively. Complete motif coverage with ensemble approach results for MEME-MAST tool can be referred in **Supplementary Table S3.2**.

#### 3.6. Summary

##### 3.6.1. Summary of Main Model

###### Human Host

In our main model, we have used 845 IL4-inducing, MHC class-II binders as positive dataset while for negative data we combine 845 IL4 non-inducing, MHC class-II binders as well as non-binders. Both positive and negative datasets are completely different as we preprocess the data to remove any redundancy. This dataset is then used for classification purpose. We have applied different approaches such as alignment-based approach (BLAST & Motif) and alignment free approach (ML, DL, LLM) to classify between IL4-inducing and non-inducing peptides. Using the alignment-based approach “BLAST”, we got 126 hits which covers 54 sequences out of 338 sequences of validation set. For motif search we used MERCI software, here we obtained 5 motifs using default parameters which covers only 8 sequences out which 1 sequence got incorrect hit, while using MEME MAST we got 14 motifs covering 38 sequences with 8 incorrect hits. Then we move forward to utilize, alignment free approaches. Firstly, we extracted 9189 composition-based features such as amino acid composition (AAC), dipeptide composition (DPC) and tripeptide composition (TPC) features. Then we applied different ML-based classifiers on composition-based features, in this task we found that extra tree classifier outperforms over other and achieved a highest AUC of 0.76 on AAC and DPC features. Then we applied different feature selection techniques such as svc-l1, mrmr, sfs, and mean based univariate analysis on all composition-based features. In this process, we obtained highest AUC of 0.80 on top 300 features selected using mean based univariate feature selection technique. We have also applied DL based classifiers such as CNN and TabNET on composition-based features and achieved AUC of 0.66 using CNN classifier on RRI feature. After that, we have also applied large language protbert model, where we achieved highest AUC of 0.72 on finetune model and AUC 0.75 on embeddings extracted using finetune model.

In last, we have developed hybrid models by combining alignment based and alignment free approaches. By combining BLAST hits with best ML scores, we achieved an AUC 0.76, by combining MERCI hits with best ML scores, we obtained an AUC of 0.80 while using MEME MAST hits with ML scores, we achieved AUC of 0.70. Here, our hybrid approach using motif hits by MERCI and best ML scores achieved highest AUC 0.80 which is similar to our best ML performance i.e; AUC of 0.80 on top 300 features selected using mean based univariate feature selection technique. We have incorporated our best model in the webserver, standalone, github package and pypi package.

###### Mouse Host

In our main model, we have used 560 IL4-inducing, MHC class-II binders as positive dataset while for negative data we combine 560 IL4 non-inducing, MHC class-II binders as well as non-binders. Both positive and negative datasets are completely different as we preprocess the data to remove any redundancy. This dataset is then used for classification purpose. We have applied different approaches such as alignment-based approach (BLAST & Motif) and alignment free approach (ML, DL, LLM) to classify between IL4-inducing and non-inducing peptides. Using the alignment-based approach “BLAST”, we got 18 hits which covers 11 sequences out of 224 sequences of validation set. For motif search we used MERCI software, here we obtained 1 motif using default parameters which covers only 1 sequence, while using MEME MAST we got 4 motifs covering 9 sequences with 3 incorrect hits. Then we move forward to utilize, alignment free approaches. Firstly, we extracted 9189 composition-based features such as amino acid composition (AAC), dipeptide composition (DPC) and tripeptide composition (TPC) features. Then we applied different ML-based classifiers on composition-based features, in this task we found that random forest classifier outperforms over other and achieved a highest AUC of 0.70 on DPC feature. Then we applied different feature selection techniques such as svc-l1, mrmr, sfs, and mean based univariate analysis on all composition-based features. In this process, we obtained highest AUC of 0.82 on top 300 features selected using mean based univariate feature selection technique. We have also applied DL based classifiers such as CNN and TabNET on composition-based features and achieved AUC of 0.64 using CNN classifier on TPC feature. After that, we have also applied large language protbert model, where we achieved highest AUC of 0.66 on finetune model as well as on embeddings extracted using finetune model.

In last, we have developed hybrid models by combining alignment-based and alignment-free approaches. By combining BLAST hits with the best ML scores, we achieved an AUC of 0.71; by combining MERCI hits with the best ML scores, we obtained an AUC of 0.80. While using MEME MAST hits with ML scores, we achieved an AUC of 0.70. Here, our hybrid approach using motif hits by MERCI and best ML scores achieved the highest AUC of 0.80, which is less than our best ML performance, i.e., an AUC of 0.82 on the top 300 features selected using the mean-based univariate feature selection technique. We have incorporated our best model in the webserver, standalone, GitHub package and Pypi package.

##### 3.6.2. Summary of Alternate1 and Alternate2 Models

###### Human Host

The alternate1 & alternate2 model, we have used 845 IL4-inducing, MHC class-II binder peptides as positive data while 516 IL4 non-inducing, MHC class-II binder peptides as negative data for alternate1 and 845 IL4 non-inducing, MHC class-II non binder peptides as negative data. Both positive and negative datasets are completely different as we preprocess the data to remove any redundancy. We follow the same workflow on alternate1 and alternate2 models for classification which we follow in main model.

In alternate1 model, our best model was developed using DPC feature that obtained highest AUC 0.82 using extra tree classifier on validation set. While in alternate2 model, highest AUC 0.85 was achieved on validation set using top 300 features selected using mean based univariate analysis.

###### Mouse Host

The alternate1 & alternate2 model, we have used 560 IL4-inducing, MHC class-II binder peptides as positive data while 560 IL4 non-inducing, MHC class-II binder peptides as negative data for alternate1 and 560 IL4 non-inducing, MHC class-II non binder peptides as negative data. Both positive and negative datasets are completely different as we preprocess the data to remove any redundancy. We follow the same workflow on alternate1 and alternate2 models for classification which we follow in main model.

In alternate1 model, our best model was developed using top 300 features selected using mean based univariate analysis that obtained highest AUC 0.82 using logistic regression classifier on validation set. While in alternate2 model, AAC feature achieved highest AUC 0.88 on validation set using random forest classifier on validation set. We have incorporated our best models in webserver, standalone, github package and pypi package.

## Web Server

In order to serve the scientific community, we provided an easy-to-use web-based interface to our users (http://webs.iiitd.edu.in/raghava/il4pred2/). IL4Pred2 web server would aid our fellow researchers, to easily scan and predict the IL4 inducing peptides. This online tool has been designed using a responsive HTML template which facilitates web page compatibility among various devices such as mobile, tablets and desktops. This web interface comprises three major modules known as: “Predict”, “Design”, and “Protein Scan”. Standalone version of this tool can be accessed using “https://webs.iiitd.edu.in/raghava/il4pred2/download.php” and is also available at GitHub “https://github.com/raghavagps/il4pred2”.

### Benchmarking with exiting methods

It is essential to compare recently developed approaches with those that already exist to assess the application and limitation of our method. In past, researchers made efforts to predict IL-4 inducing peptides. The first method was developed by Dhanda et al, in 2013 to predict IL-4 inducing peptides and achieved a maximum accuracy of 75.76% and MCC of 0.51 by utilizing amino acid pairs and motif information [22]. After that in 2023, Hassan et al, developed another method named “Meta-IL4” based on ensemble approach. This method utilized amphiphilic pseudo-amino acid composition (APAAC) feature to predict IL-4 inducing peptides and reported highest accuracy of 90.70%. Recently, a new method named as “Plm-Il4” has been developed by Liu et al, using ESM-2 based model achieved accuracy of 93% [23]. The major limitation of all the methods is that they have been developed using experimental IL-4 inducing data irrespective of their host. To overcome this limitation, we have developed a host specific method to classify IL4-inducing peptides using human and mouse as host. In the present study, the compositional analysis of human and mouse host shown a major difference in average amino acid composition. The amino acid residue leucine is shown to be higher average composition in Human host while glycine and proline has higher average amino acid composition in Mouse host as shown in **Figure 4.2**.

We have performed a benchmarking study of IL4pred-2 method using validation dataset and newly added data, which is not present in previous methods, to calculate the performance over other methods. Here, we have observed that the main validation dataset of Human and Mouse host on hybrid model (SVM and motif) of IL4pred achieved an accuracy of 75% and 79% on both human and mouse data while it obtained an accuracy of 50% and 63% using newly added data on human and mouse. The Plm-Il4 achieved accuracy of 61% and 66% on validation data of human and mouse and accuracy of 64% and 81% on newly added data of human and mouse (**see Table 11.1**). The above defined models performed well on validation data of IL4pred2 as these methods already trained on around 60% same positive and 51% same negative dataset of IL4pred2.

**Table 11.1:**
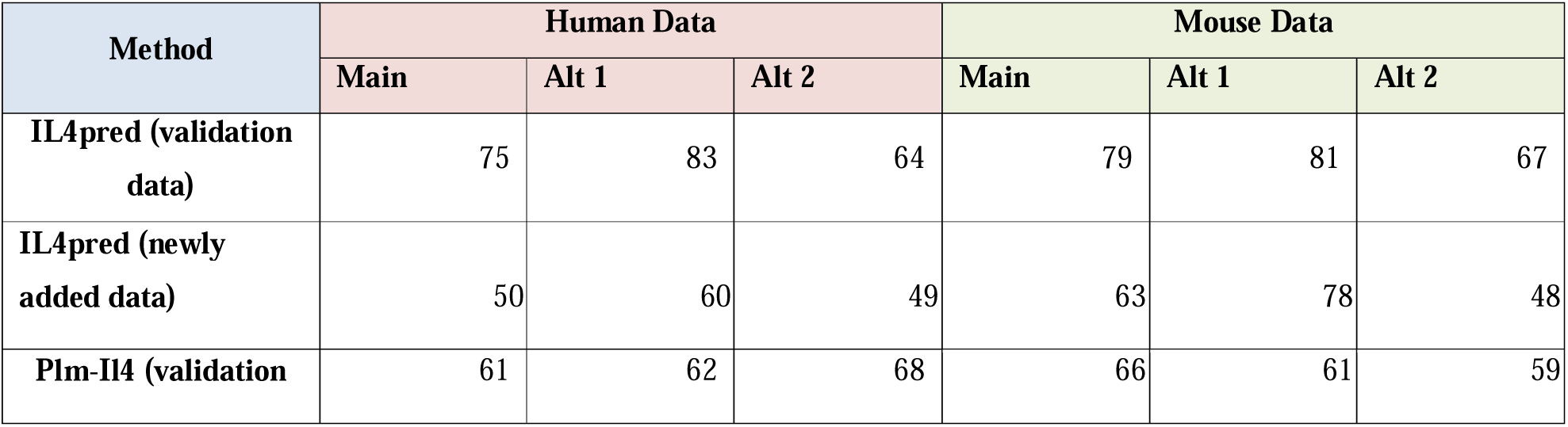
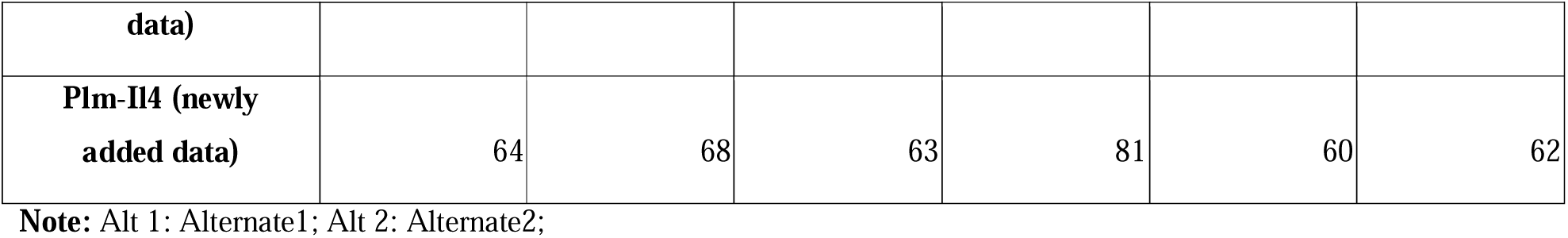
This table depicts the benchmarking accuracy of validation data of IL4pred2 over different methods.

We have also performed a reverse analysis of IL4pred-2 human model performance evaluation on IL4pred-2 mouse model and vice versa. This analysis was performed to validate our hypothesis that human model not working well on mouse data and mouse model does not work well on human data due to their different amino acid composition. Here, we have observed that main validation dataset of human achieved only 0.50 AUC using main model of mouse host. It can be clearly seen that mouse model is unable to discriminate between il4-inducing data of human host. Then we check performance of mouse main validation data on human main model and achieved an AUC of 0.58 (**see Table 11.2**). These finding clearly support our hypothesis about different models for human and mouse hosts. Previously, we had already shown that discrimination between hosts in another method which is designed for discovering interferon-gamma inducing peptides in human and mouse [45]. The above explained results are shown in **Table 11**, for detailed results please refer **Supplementary Table S10**.

**Table 11.2:** This table depicts the benchmarking performance of validation data of IL4pred2 over different methods.

| Mouse Model | Human Independent Data |
| --- | --- |
| Main AUC | 0.50 |
| Alternate1 AUC | 0.49 |
| Alternate2 AUC | 0.62 |
| Human Model | Mouse Independent Data |
| Main AUC | 0.58 |
| Alternate1 AUC | 0.52 |
| Alternate2 AUC | 0.60 |

## Discussion

In the era of peptide therapeutics, IL4 based drugs and therapeutics plays a pivotal role in disease prevention. As it originates from T helper 2 cells along with other cytokines, it helps in inhibiting T_H1_-like and T_H_17-mediated inflammatory processes and act as anti-inflammatory cytokine. It also plays important role in tissue regeneration and the host’s defense against parasites [46]. Although, it had also been found in the development of cancer, autoimmune disorders, and allergy diseases [15]. Research on IL-4’s potential in immunotherapy and disease modulation is continuing, and given its wide immunological impact, it continues to be a prospective therapeutic target for several diseases [47]. Abrocitinib, Dupilumab, Upadacitinib, Tofacitinib are the few U.S. Food and Drug Administration, and the European Medicines Agency approved therapies which targets IL-4/IL-13 signaling. These drugs targets IL-4Rα receptor, and/or molecules of JAK1, JAK3 and used for treatment purposes of atopic dermatitis, severe asthma, chronic sinusitis, nasal polyps and rheumatoid arthritis [15].

In this study, we tried to develop a host specific prediction method where we used two hosts: Human and Mouse. We have gathered experimentally validated MHC-class II binding IL-4 inducing peptides of Human and Mouse host for prediction task. Each host method consists of three different models depending on nature of IL4 non-inducing peptides. Firstly, we have performed preliminary analysis using main dataset of human and mouse hosts. The positional and compositional analysis amino acid residues shows that leucine is abundant in human host while glycine and proline are abundant in mouse host. We have also implemented alignment-based approaches such as BLAST and Motif (MERCI and MEME) and alignment free approaches such as machine learning based classifiers, deep learning-based classifiers and large language models for classification and prediction of IL-4 inducing peptides. The classification task is completely dependent on most relevant feature. To extract important features, we have performed different feature selection techniques such as SVC-L1, mRMR, SFS, LR-based and univariate (mean-based) feature selection method. As we have prepared three different datasets for each host, different features were used to develop models on individual datasets. Human and Mouse main model is developed using top 300 composition-based selected features from univariate feature selection method. We have obtained 0.80 AUC with 0.45 MCC on human main model while 0.82 AUC with 0.50 MCC on mouse main model. The best models of all the three datasets for both hosts are incorporated in webserver and standalone packages.

Most of the existing studies utilized single dataset for developing prediction model for this type of studies. This raised an obvious question, why we used three types of datasets for building models. In all three datasets, positive dataset is identical that contain IL-4 inducing peptides extracted from IEDB. The major challenge is to choose a negative dataset which is acceptable by the scientific community. In Alterante1 dataset, we have considered MHC class II binders which don’t induce IL-4 as the negative set. This dataset contains experimentally confirmed IL-4 inducing and IL-4 non-inducing MHC class II binding peptides. The majority of existing studies generally use this type of dataset for model building. Model built on the Alternate1 dataset is intended to differentiate between IL-4-inducing and IL-4 non-inducing MHC Class II binders. This model is most helpful if the user knows that their peptide is an MHC Class II binder, but it will fail if peptide is non-binder. To offer different solutions, we have introduced Alternate2 dataset that contain non-MHC Class II binders as the negative set. Model built on this dataset is useful for separating IL-4 inducing peptides from non-MHC Class II binders. But can’t differentiate between MHC Class II binders which do or don’t induce IL-4. In real-world, most of time user have no idea whether given peptide an MHC Class II binder or not. Then model built on above datasets will be failed. To overcome above drawbacks, we have proposed a dataset called Main dataset. This dataset contains both MHC binders and non-binders as negative dataset, all of which are experimentally confirmed IL-4 non-inducing peptides. This model is most appropriate for scenarios in which the user does not have any information regarding whether their peptide is an MHC binder or not. To sum up, we trained our models on three different datasets to cater to different ranges of user requirements under different scenarios.

## Conclusion

In this study, we have attempted a systemic approach for benchmarking the existing methods for scanning and predicting the IL4 inducing peptides available in public domain. The aim of this study was to expedite the time-consuming process and higher precision involved in designing novel drug adjuvant and immunotherapies revolving around IL4 cytokine. Although generating an immune response via inducing a cytokine/ interleukin in disease patients is a complex and challenging problem, we hope that this tool would aid in identifying peptide-based therapeutics against various disease states.

## Acknowledgments

Authors are thankful to the Department of Science and Technology (DST-INSPIRE), University Grants Commission (UGC), Department of Biotechnology (DBT), and the Department of Computational Biology, IIITD New Delhi for fellowships, financial support, infrastructure and facilities. We would like to acknowledge that Figures were created using BioRender.com.

## Data Availability Statement

All the datasets used in this study are available at the “il4pred2” webserver, https://webs.iiitd.edu.in/raghava/il4pred2/download.html.

## Preprint

The preprint is available at biorxiv https://doi.org/10.1101/2025.04.23.650150.

## Author Contributions

RT and GPSR collected and processed the datasets. RT, NM and GPSR implemented the algorithms and developed the prediction models. RT, NM and GPSR analyzed the results. RT created the front-end user interface and back end of the webserver. RT, SJ and GPSR penned the manuscript. SM prepared the figures and pypi package for the manuscript. RT, NM, SM, SJ and GPSR reviewed the manuscript. GPSR conceived and coordinated the project. All authors read and approved the final manuscript.

