## Supplementary figures and images for "IL4Pred2: Prediction of Interleukin-4 Inducing Peptides in Human and Mouse"

### Supplementary Figure S1

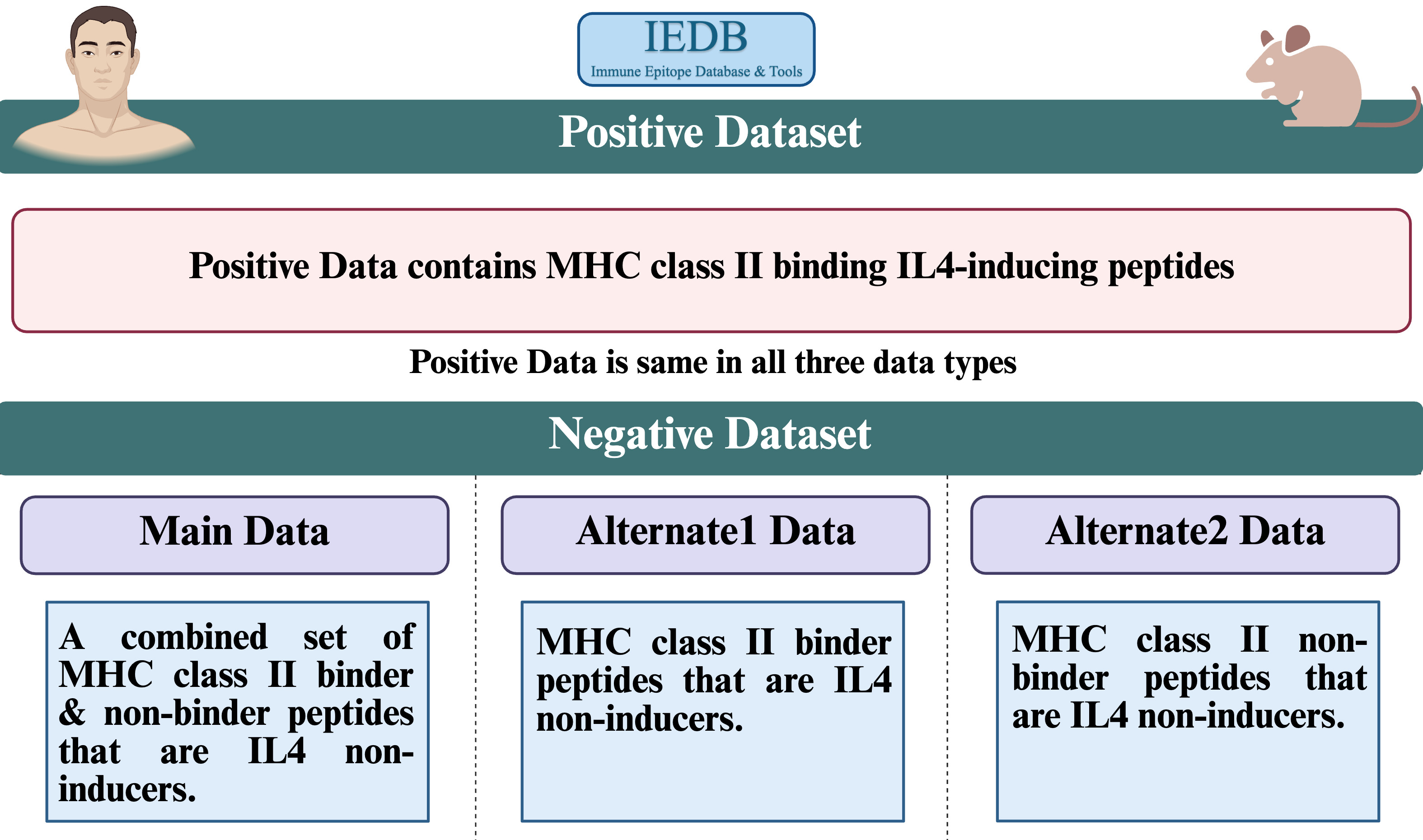
